# Stability of neural representations in the auditory midbrain across the lifespan despite age-related brainstem delays

**DOI:** 10.1101/2022.11.20.517243

**Authors:** Rüdiger Land, Andrej Kral

**Affiliations:** Institute for Audioneurotechnology, Department of Experimental Otology, Hannover Medical School, Germany; Australian Hearing Hub, School of Medicine and Health Sciences, Macquarie University, Sydney, Australia

**Keywords:** brainstem, auditory midbrain, gap detection, temporal resolution, auditory temporal processing, ABR, auditory brainstem response, aging, neural representation

## Abstract

The extent to which aging of the central auditory pathway impairs auditory perception in the elderly independent of peripheral cochlear decline is debated. To cause auditory deficits in normal hearing elderly, central aging needs to degrade neural sound representations at some point along the auditory pathway. However, inaccessible to psychophysical methods, the level of the auditory pathway at which aging starts to effectively degrade neural sound representations remains poorly differentiated. Here we tested how potential age-related changes in the auditory brainstem affect the stability of spatiotemporal multiunit complex speech-like sound representations in the auditory midbrain of old normal hearing CBA/J mice. Although brainstem conduction speed slowed down in old mice, the change was limited to the sub-millisecond range and only minimally affected temporal processing in the midbrain (i.e. gaps-in-noise sensitivity). Importantly, besides the small delay, multiunit complex temporal sound representations in the auditory midbrain did not differ between young and old mice. This shows that although small age-related neural effects in simple sound parameters in the lower brainstem may be present in aging they do not effectively deteriorate complex neural population representations at the level of the auditory midbrain when peripheral hearing remains normal. This result challenges the widespread belief of ‘pure’ central auditory decline as an automatic consequence of aging. However, the stability of midbrain processing in aging emphasizes the role of undetected ‘hidden’ peripheral damage and accumulating effects in higher cortical auditory-cognitive processing explaining perception deficits in ‘normal hearing’ elderly.

## 1. Introduction

Problems in speech understanding are common in the elderly. Although these perceptual problems are dominantly caused by peripheral age-related cochlear damage, aging of the central auditory pathway itself may additionally deteriorate auditory neural signals (Pichora-Fuller and Singh, 2006). Understanding the extent to which central aging contributes to auditory deficits - independent of peripheral damage - is important to explain self-reported hearing problems in normal hearing elderly, and to maximize the user benefits of hearing aids (Gates, 2012; Davidson et al., 2021; Lentz et al., 2022).

Three physiological causes can in principle lead to age-related deficits in auditory perception: cochlear, central, and cognitive decline (Humes and Dubno, 2010; Humes et al., 2012; Sardone et al., 2019; Lentz et al., 2022). By far the most prominent, and best understood factor is age-related *peripheral* cochlear damage affecting the *input level* of acoustic signals (Gates and Mills, 2005; Keithley, 2020; Wang and Puel, 2020; Eckert et al., 2021). The second factor, recently receiving more attention, lies in the age-related decline of cognitive processing of auditory information, such as attention, or working memory (Pichora-Fuller and Singh, 2006; Gordon-Salant et al., 2020; Humes et al., 2022; Lentz et al., 2022). The third factor is central aging, defined as the age-related deterioration of the *transmission of information* throughout the central auditory pathway (Walton, 2010; Humes et al., 2012; Atcherson et al., 2015; Ouda et al., 2015; Anderson and Karawani, 2020), and can be anatomically situated between cochlea and higher auditory-cognitive-association structures.

In comparison to peripheral and cognitive effects (Eckert et al., 2021), age-related *central* processing deficits - sometimes referred to as ‘central presbycusis’-are less clear. Mixed evidence has led some authors to question the existence of a ‘pure’ age-related central auditory decline in isolation without being confounded by ‘indirect’ effects of cochlear damage or top-down cognitive factors (Humes et al., 2012; Atcherson et al., 2015; Land and Kral, 2022). This is reflected by the fact, that variability in findings on simple measures support both sides, the preservation of central temporal acuity in aging, as well as its decline, e.g. in gap detection, envelope detection, or transmission speed/slowing (Shen, 2014; Quraishe et al., 2020). Even with the presence of small effects on basic parameters of central processing in aging, for ‘true’ central presbycusis, however, it needs to be shown that age-related central decline degrades neural representations of complex speech sounds at some level of the auditory pathway to effectively impact speech perception – independent of peripheral cochlear damage and cognitive decline (Humes et al., 2012). The ‘where’ and ‘extent’ of such central presbycusis on the population neuronal level remain poorly differentiated, as they cannot be addressed with psychophysical methods.

We here asked whether central aging can effectively deteriorate neural representations of complex signals on the level of the auditory midbrain in absence of age-related peripheral hearing loss. In addition, we tested how strongly this is affected by the potential presence of age-related changes on the level of the lower auditory brainstem.

To study this, we measured the extent of age-related changes in central conduction speed and temporal resolution in the brainstem and compared it to the age-stability of neural ‘speech-like’ sound representations in the auditory midbrain of normal hearing CBA/J mice. Specifically, we combined measures of ABR interpeak intervals, and ABR gap detection in the brainstem, with recordings of multiunit population activity in the IC to gaps-in-noise and complex ‘speech-like’ sounds. We then compared central brainstem transmission speed, temporal resolution, and neural representations of complex stimuli between young and aged mice up to 24 months old, that were selected for normal hearing to prevent influence from peripheral damage confounds. Here we found that although small age-related delays may occur in the lower auditory brainstem, neural sound representations remain stable in the auditory midbrain of old mice.

## 2. Material and Methods

### 2.1. Animals

Female CBA/J mice in an age range between 3 to 24 months were tested (n = 25) and had been selected to have normal peripheral hearing. CBA/J can have robust hearing thresholds throughout life, with no or only relatively mild hearing loss in aging (Willott et al., 1988; Li and Borg, 1991; May et al., 2006; Sha et al., 2008; Ohlemiller et al., 2010). The age range of the CBA/J mice used can be translated approximately to a human age range from 5 to 80 years (Dutta and Sengupta, 2016). Experiments were conducted in accordance with ethical standards for the care and use of animals in research and the German law for the protection of animals. The experiments were approved by the ethics committee of the state of Lower Saxony. All animals were handled and housed according to German (TierSchG, BGBl. I S. 1206, 1313) and European Union (ETS 123; Directive 2010/63/EU) guidelines for animal research and the described animal experiments were approved by the German state authorities (Lower Saxony State Office for Consumer Protection and Food Safety [LAVES] and monitored by the university animal welfare officer.

### 2.2. General Anesthesia

Animals were anesthetized by an initial dose of intraperitoneal injection of 100mg/kg Ketamine (Ketamin Graeub, Albrecht GmbH, Germany); and 4mg/kg Xylazine (Albrecht GmbH, Germany). Animals were placed on a temperature probe-controlled heating pad (TC-1000 Temperature Controller, CWE Inc., USA) with a rectal probe for measuring and keeping core temperature in a range of 37.6–37.8 degrees Celsius. For ECG monitoring, silver wire electrodes were used to monitor ECG and heart rate during the whole experiment. At initiation of anesthesia and during the experiment the toe-pinch reflex was assessed every 15 min to ensure adequate anesthesia depth. If needed a second dose of Ketamine/Xylazine was applied to ensure adequate anesthesia depth, usually after 45-60 minutes after the first dose.

### 2.3. ABR recordings

The auditory brainstem responses (ABRs) were recorded with teflon coated subdermal monopolar needle electrodes (0.35 mm, 15 mm, GVB Gelimed, Germany) positioned at the vertex against a reference behind the right ear and a ground in the neck or the other ear as previously described (Willott, 2006; Land et al., 2016). ABR signals were amplified 10,000 times with a custom-built amplifier (Otoconsult Co, Frankfurt, Germany) and were wideband filtered 1-9000 Hz and subsequently digitized and recorded with a Digital Lynx SX recording system (Neuralynx, Bozeman, USA) at a 32 kHz sampling rate with 24 bit A/D converter resolution. Signals were recorded with Cheetah software 5.11. saved on a Z800 HP computer and stored in Neuralynx.nev file format.

### 2.4. Multielectrode array recordings

Multiunit activity in the IC was measured with a 32-channel linear multisite electrode array (1 × 32, site distance 100 μm, site area 177 mm^2^, Impedance 2-3 MOhm at 1 kHz, NeuroNexus, USA) against a frontal epidural silver-ball electrode, which served as reference. After trepanation of the overlying skull, the electrode was inserted vertically into the middle portion of the visible IC surface until it spanned the full IC and covered a depth down to approximately 3000 μm. This resulted in penetration with the lower electrode contacts below and the topmost electrodes above the surface of the IC. The electrode was cleaned after each experiment with NaCl (B. Braun, Melsungen, Germany) and contact lens solution (Progent, Menicon) and reused multiple times. The neuronal signals were recorded with Neuralynx Digital Lynx SX (Neuralynx, Bozeman, USA) and preamplified with a Neuralynx HS36 headstage (Neuralynx, Bozeman, USA), referenced to a frontal silver ball electrode. The NeuroNexus 1 × 32 electrode was attached to the headstage with a Neuronexus probe connector (ADPT-HS36-N2T-32A, Neuralynx, USA). Signals were filtered during the recording at 0.1 Hz −9000 Hz and digitized at 32 kHz with 24-bit A/D converter resolution. Signals were recorded with Cheetah software 5.11. saved on a Z800 HP computer and stored in Neuralynx file format.

### 2.5. Acoustic Stimulation

The mice were placed in a dark sound proof chamber (IAC acoustics, Winchester, UK) during the experiment. The acoustic stimuli were presented in free-field with a tweeter speaker (Vifa XT300 K/4) positioned 10 cm in front of the animal at 0 cm elevation. Acoustic stimuli were generated with Matlab (2018b, The Mathworks) running on a stimulation PC (Dell Precision T5810) and output via a multifunction I/O device (National Instruments, PCIe-6353 with a BNC 2090A). Analog signals were then sent to an attenuator then passed on to the free-field tweeter speaker.

#### 2.5.1 Click thresholds

Alternating condensation and rarefaction clicks of 5 μs duration were presented at levels from 20 dB to 95 dB peSPL in 5 dB steps. The inter-click interval was 100 ms and clicks were repeated 600 times. Click level was defined as the sinusoidal relative level and calculated for the rectified amplitude of the response, as measured with a calibrated Bruel and Kjaer microphone (Free-field 1/4 Microphone Type 4939, Bruel and Kjaer, Denmark). To measure the click level, the microphone was placed at the same position as the mouse head during the experiment at the same distance as the speaker.

#### 2.5.2 Gaps-in-noise

Gaps were presented in white noise at five gap intervals from 0, 0.5, 1, 5, and 10 milliseconds. The stimulus consisted of 500 ms white noise with gaps of different durations inserted after 250 ms. White noise onset and intermitted gaps were not ramped and presented at 90 dB SPL. Each condition was repeated 300 times. White noise distribution was randomly generated for each condition.

#### 2.5.3 Complex sounds

Two types of complex sounds were used in this study. The first complex sound used was an electronic version (Tetris/Hardcore remix – Lumin8) of the Russian folk song *Коробе́йники* (Korobéyniki), better known as the Tetris theme song, sampled at 44.1 kHz and presented free-field at average 85 dB SPL. The duration of the song snipped was 56 seconds and was repeated 5 times. The second type of complex sounds were 24 words with different consonants presented by a female speaker (International Phonetic Association, IPA Handbook Downloads), sampled at 20 kHz and repeated 20 times presented free-field at an average of 85 dB SPL.

### 2.6 Data pre-processing

#### 2.6.1 ABR pre-processing

The Neuralynx data files were imported and processed with Matlab (Matlab2018b, The Mathworks). For the ABR signal analysis, the digitized analog signals were filtered in a bandwidth of 300-3000 Hz (narrow-band) and 30-3000Hz (wide-band) with a zero-phase 10^th^ order Butterworth digital filter (matlab function *filtfilt, butter*). We analyzed the ABR with a traditional narrow filter (300-3000 Hz) and one with wider filter settings (30-3000Hz), which allowed to visualize the IC-related P_0_ component (Land et al., 2016; Duque et al., 2018). After filtering, the ABR signal was down-sampled to a 30 kHz sampling rate using the Matlab function resample. The ABR signal was then segmented into trials and averaged over the trials. For the noise onset responses, the first three ABR waves were always best discernible, with waves IV and V sometimes less prominent. We used the nomenclature with Roman numerals from I to V with an additional later peak P_0_ representing IC activity (Land et al., 2016; Duque et al., 2018).

#### 2.6.2 MUA pre-processing

For the multiunit analysis, the digitized analog raw signal was filtered with the Matlab function filtfilt for zero-phase digital filtering with a bandpass of 500–9000 Hz. We used a 10^th^-order Butterworth bandpass filter generated with the Matlab function *butter*. A fixed threshold of 3.5 SD of the background activity was then used to collect time points of all spiking activity above that threshold that were then defined as multiunit activity.

### 2.7 Data analysis

#### 2.7.1 ABR analysis

##### Click thresholds

Individual click ABR thresholds were defined as the dB SPL level, at which the root-mean-square (RMS) level of the ABR response (1-10 ms post-stimulus interval) was first significantly different from the baseline activity. The automatic threshold detection was independently visually verified.

##### ABR wave latencies

Peak latencies were determined for the ABR onset response and ABR gap response at 10 ms. The ABR wave latencies were determined as peak latencies of the respective waves. Interpeak intervals were derived by subtracting the respective peak latencies. We determined peak latencies of wave I-V in addition to wave P_0_, previously described as an IC component in the mouse ABR (Land et al., 2016).

##### Gap ABR spectral analysis

Multi-taper spectrograms were calculated using the Chronux Toolbox (http://chronux.org/) for Matlab (Bokil et al., 2010). For each mouse and gap duration, multi-taper spectrograms were determined for 13 ms intervals of the ABR response in a 6 ms window and 1 ms step-size and tapers with a time-bandwidth product TW=2 and a number of tapers K=3. Spectrograms of baseline activity were subtracted from the ABR spectrograms and the result was then normalized to the maximum. Center frequencies were then determined by calculating the weighted centroid using the Matlab *regionprops* function.

##### Gap ABR

To determine the gap detection threshold, we calculated the ABR gap response. This was done by determining the peak-to-peak amplitude of the evoked ABR response after the gap. The average ABR response was calculated from the 300 repetitions for each condition. Two metrics were used to quantify ABR gap response. The peak-to-peak amplitude of the ABR response and the RMS (root-mean-square) value of the ABR response. These were A) calculated as absolute values with subtracted baseline, and B) calculated as relative to the onset ABR of the noise burst (recovery function).

##### Multiunit gap responses and gap sensitivity

We pooled the five most responsive sites in each mouse for each IC recording and analyzed the multiunit gap response as the mean spike count in a 15 ms window after the gap offset. The presence of significant gap responses (gap sensitivity) was determined by a permutation test. We generated a surrogate baseline distribution by randomly selecting spike counts in 15 ms windows of multiunit activity previous to the gap (400 permutations). We then defined significant responses as responses exceeding a threshold level of the 10 % highest spike counts of the baseline distribution.

##### Correlation of neural sound representations between mice

We defined the neural sound representation as the averaged spatial multiunit responses of the electrode contact with the maximum response and the 3 adjacent electrode contacts above and below. Responses were then averaged over trial repetitions of the respective sounds. The pairwise crosscorrelation of multiunit responses between all mice was then calculated with the Matlab function *xcorr*. The cross-correlation was normalized so that the auto-correlations at zero lag equal 1. The peak of the normalized cross-correlation function was used to determine the shift/delay of the multiunit response in old mice in comparison to young mice. The correlation strength was defined as the peak of the cross-correlation. The effect of age on neural sound representations was then tested by comparing the mean pairwise correlation between the respective age groups.

##### Correlation of neural sound representations with complex sounds

We calculated the crosscorrelation between the down-sampled speech signals and the multiunit responses for each mouse as a measure of sound representation of complex signals. The neural sound representation was defined as the averaged spatial multiunit responses of the electrode contact with the maximum response and the 3 adjacent electrode contacts above and below. The neural sound representation was correlated to the 3-10 kHz band power envelope estimate. This was derived from the multi-taper spectrogram with 5 ms window and 1 ms step size and tapers with a time-bandwidth product TW=1 and number of tapers K=2. The envelope signal was then downsampled to 1000 Hz, and the cross-correlation between neural response and envelope signal was calculated.

### 2.8 Statistical methods

Statistical tests are described in the text, where applied. In summary, we used ANOVA and two-way ANOVA (with the Matlab functions *anova1* and *nanovan*), and the significance level was 5 %. Measures of variance and error bars are standard deviation (SD) or standard error of the mean (SEM) and are denoted at respective parts in the text and figure legends. Bootstrapped confidence intervals were calculated using the Matlab function *bootci*. A linear regression model was fitted to the data with the Matlab function *fitlm*, results and fit model functions are provided in Table 1 and Table 2.

**Table 1.** Linear regression model of ABR peak latency change with age (ms/day)

| ABR Wave | Linear model | R<sup>2</sup> | F-Statistic | p-value |
| --- | --- | --- | --- | --- |
| I | $-0.0003x + 1.26$ | 0.24 | $F(1,22) = 6.85$ | 0.016 |
| II | $0.0001x + 2.05$ | 0.05 | $F(1,22) = 1.26$ | 0.275 |
| III | $0.0003x + 2.64$ | 0.35 | $F(1,22) = 12.0$ | 0.002 |
| IV | $0.0005x + 3.42$ | 0.27 | $F(1,22) = 7.43$ | 0.013 |
| V | $0.0009x + 4.63$ | 0.30 | $F(1,22) = 10.1$ | 0.004 |
| $P_0$ | $0.0027x + 6.90$ | 0.31 | $F(1,22) = 10.4$ | 0.003 |

**Table 2.** Linear regression model for ABR interpeak interval change with age (ms/day)

| Interpeak Interval | Linear model | R<sup>2</sup> | F-Statistic | p-value |
| --- | --- | --- | --- | --- |
| **Onset** |  |  |  |  |
| I-V | $0.001x+3.16$ | 0.40 | $F(1,22) = 14.6$ | 0.0009 |
| I-P<sub>0</sub> | $0.003x+5.75$ | 0.37 | $F(1,22) = 13.1$ | 0.015 |
| **10ms Gap** |  |  |  |  |
| I-V | $0.0004x+3.57$ | 0.12 | $F(1,22) = 1.95$ | 0.059 |
| I-P<sub>0</sub> | $0.0009x+6.87$ | 0.14 | $F(1,22) = 3.75$ | 0.067 |

## 3. Results

### 3.1 Auditory brainstem responses and IC multiunit recordings in young and old CBA/J mice with normal hearing

We measured auditory brainstem responses (ABR) to gaps-in-noise in CBA/J mice in an age range from 2 to 24 months **(Figure 1A**). Mice were selected to have essentially normal click ABR thresholds < 40 dB peSPL to minimize age-related confounds by cochlear peripheral hearing loss **(Figure 1B**). The sound onset of white noise elicited ABR with waves I-V and also the late IC component P_0_ when wide band-pass filtering was used **(Figure 1C**). ABR gap responses decreased in amplitude with decreasing gap durations **(Figure 1D, E)**. The absolute power of the ABR onset response did not change with age for narrow-filter ABR onset response (r(23) = 0.19, p = 0.39) and wide-filter ABR onset response (r(23) = 0.05, p = 0.82). Additional to the ABR recordings, we recorded multiunit activity in the inferior colliculus (IC) and measured the gap sensitivity of multiunit IC activity **(Figure 1F)**.

**Fig. 1.**
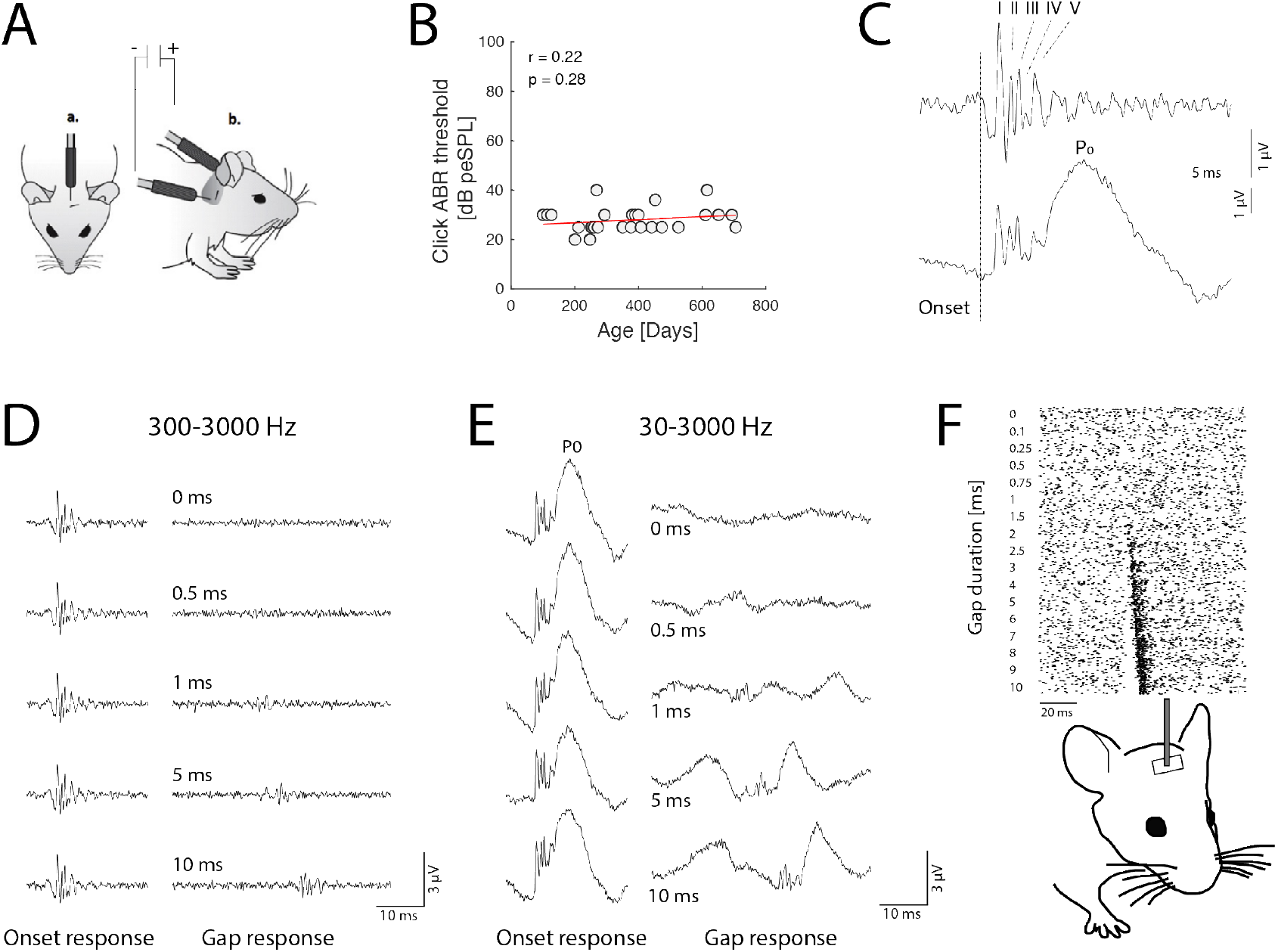
Auditory brainstem responses and IC multiunit recordings in young and old CBA/J mice with normal hearing. A) Auditory brainstem response (ABR) recording configuration with retro-auricular subcutaneous needle electrodes. B) Aging CBA/J mice were selected for normal click ABR thresholds. ABR click thresholds showed no significant correlation with age r(23) = 0.22, p = 0.28. C) ABR responses were recorded with two filter settings. The upper trace shows classical ‘narrow’ filtering 300-3000 Hz and ABR peaks I-V. The lower trace shows ‘wide’ filtering 30-3000 Hz, which visualizes the late P_0_ component, representing IC activity. D) Gap ABR from a CBA/J mouse (age: 8 months) recorded with a narrow band-pass filter. Onset responses to white noise on the left and responses to increasing gap durations from top to bottom on the right. E) Gap ABR from the same mouse as in D) with wide bandpass filter visualizing the late P_0_ component. F) Multiunit recordings from the IC of CBA/J mice with a linear multielectrode array. The raster plot shows multiunit responses from one electrode channel within the IC for decreasing gap durations (10 repetitions for each duration).

### 3.2 Age-related slowing of auditory brainstem conduction speed in normal hearing mice

Aging affected the central conduction speed in the auditory brainstem. With increasing age, ABR wave peak latencies changed significantly with exception of wave II **(Figure 2A, Table 1)**. Correspondingly, the I-V interval and the I-P_0_ interval of the onset ABR significantly increased in old normal-hearing mice **(Figure 2B, Table 2)**. The slopes for the intervals of gaps with 10 ms duration showed a similar trend, however did not fully reach a significance level of p = 0.05 **(Figure 2C, Table 2)**.

**Fig. 2.**
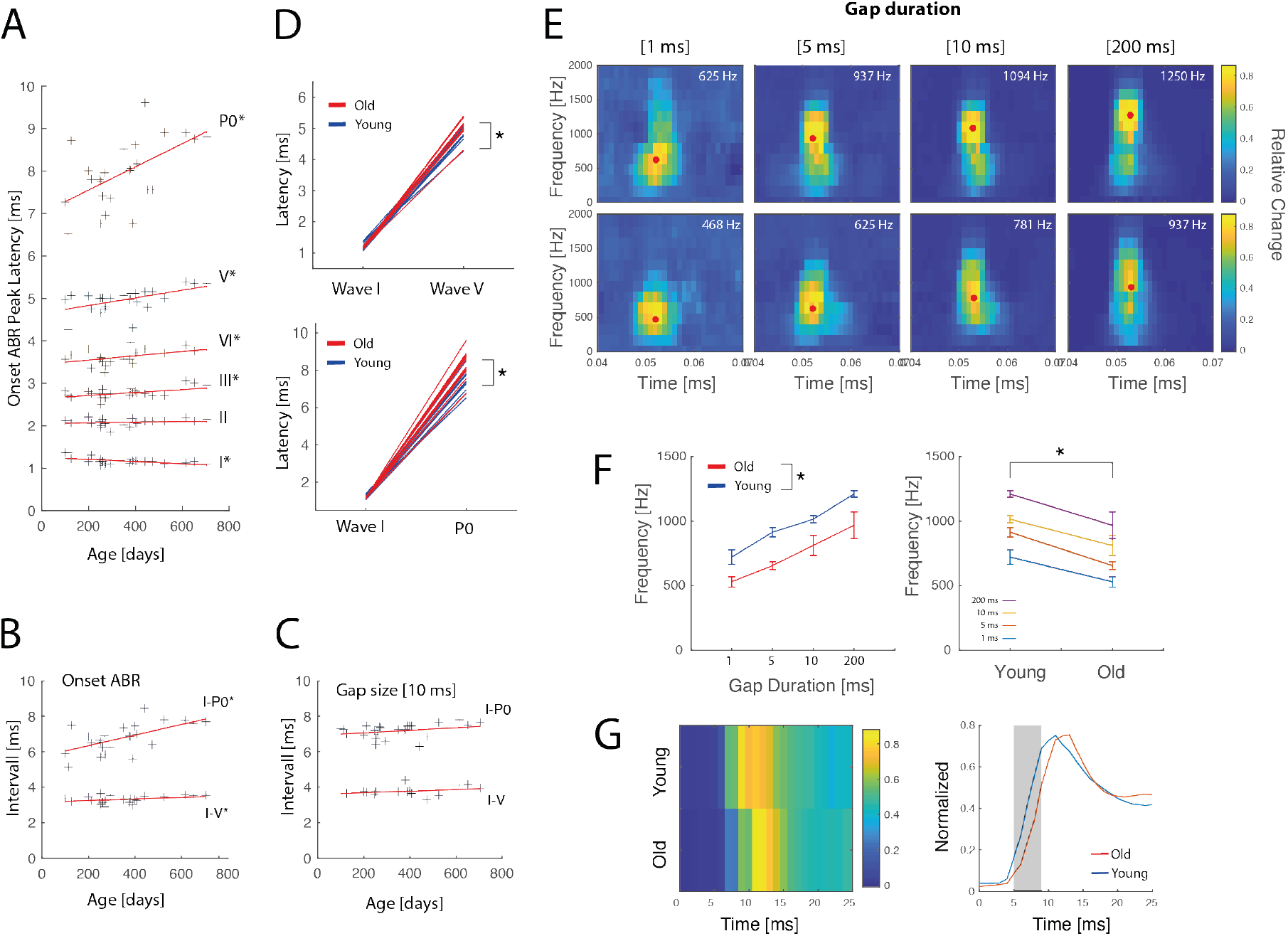
Age-related slowing of auditory brainstem conduction speed in normal hearing mice. A) Onset ABR peak latencies for waves I to V, and P_0_ over age in CBA/J mice. Red lines show linear regression for individual wave latencies. B) ABR I-V and I-P_0_ interpeak intervals for noise onset. C) ABR I-V and I-P_0_ interpeak intervals for 10 ms gap duration. D) Group comparison of wave I and V latencies (top) and I and P_0_ (bottom) latencies between old and young mice. E) Multi-taper spectrograms of gap ABRs for different durations in young (n=9; 195± 65 days; mean/SD) and old mice (n=5; 593 ± 94 days; mean/SD). F) Center frequencies of spectrograms differ in young and old mice (left) for all gap durations (right). G) Comparison of multiunit onset responses in young and old mice. Multiunit responses in old mice show a significant delay at onset in comparison to young mice (right panel, two-sample t-test, p<0.05). For visualization responses were smoothed with a 5 ms moving average filter and rescaled.

Group-wise comparison between old (> 365 days; (484(114) days mean/SD, n = 12) and young mice (< 365 days; 226(74) days mean/SD, n = 13) additionally showed differences between the onset ABR I-V interval and I-P_0_ interval **(Figure 2D)**. A two-sample t-test to compare the ABR I-V intervals between old and young mice showed no significant difference in intervals between young (4.88 ± 0.23 ms; mean/SD) and old (5.05 ± 0.29 ms; mean/SD); t(23) = −1.64, p = .11. A two-sample t-test to compare the ABR I-P_0_ intervals between old and young mice showed a significant difference in intervals between young (7.60 ± 0.64 ms; mean/SD) and old mice (8.31 ± 0.77 ms; mean/SD); t(23) = −2.49, p = .02).

Age-related slowing of central conduction time was also reflected by a decrease in the ABR spectrogram center frequency. **Figure 2E** shows averaged time-frequency representations for gap responses of 1 ms, 5 ms, 10 ms, and onset responses. The center frequency of the ABR decreased in older mice, reflecting an elongation of the ABR response (**Figure 2F)**. Twoway ANOVA showed no significant interaction of center frequency between the effects of age and gap duration (F(4,51)=1.25 p=0.3). Simple main effects analysis confirmed that gap duration had a statistically significant effect on center frequency (F(4,51)=29 p=0), and age showed a statistically significant effect on center frequency (F(1,51)=36.4 p=0). Group-wise comparison between age groups showed significant differences in center frequencies for each gap duration (**Figure 2F**, right panel, two-sample t-test, p<0.05). Last, we tested the effect of age on the latency of white noise-evoked multiunit activity in the IC. This also showed a significant onset delay in the range of 1 ms for aged animals, with additional differences in the later phase of the response (two-sample t-test with Bonferroni-Holm correction for multiple comparisons, p<0.05) (**Figure 2G**).

### 3.3 Auditory brainstem gap sensitivity remains robust over lifetime in normal hearing mice

We next tested if the age-related delays of the ABR affected ABR gap sensitivity. Gap ABRs showed no obvious change in shape in old mice **(Figure 3A)**, and the proportion of responses to decreasing gap durations did not differ between young (< 365 days; 226(74) days mean/SD, n = 13) and old (> 365 days; (484(114) days mean/SD, n = 12) mice **(Figure 3B, C**). ABR responses to gaps of 10 ms and 5 ms were present in mice of all ages, and up to 50 % of mice in both age groups still showed a gap sensitivity to 1 ms gaps. Further, gap ABR response amplitudes did not change in old mice **(Figure 3D)**. Two-way ANOVA showed no significant interaction of ABR peak-peak amplitude between age and gap duration (F(4,115)=0.59 p=0.67). Simple main effects analysis showed a statistically significant effect of gap duration on response amplitude (F(4,115)=39.10 p=0), however, age did not have a statistically significant effect on response amplitude (F(1,115)=0.01 p=0.92). The same was true for the comparison of the RMS of the ABR response between young and old mice **(Figure 3E)**. Two-way ANOVA showed no interaction of ABR RMS between age and gap duration (F(4,115)=0.3 p=0.88). Simple main effects analysis confirmed that gap duration had a statistically significant effect on response amplitude (F(4,115)=71.27 p=0), however, age did not have a statistically significant effect (F(1,115)=0.01 p=0.91).

**Fig. 3.**
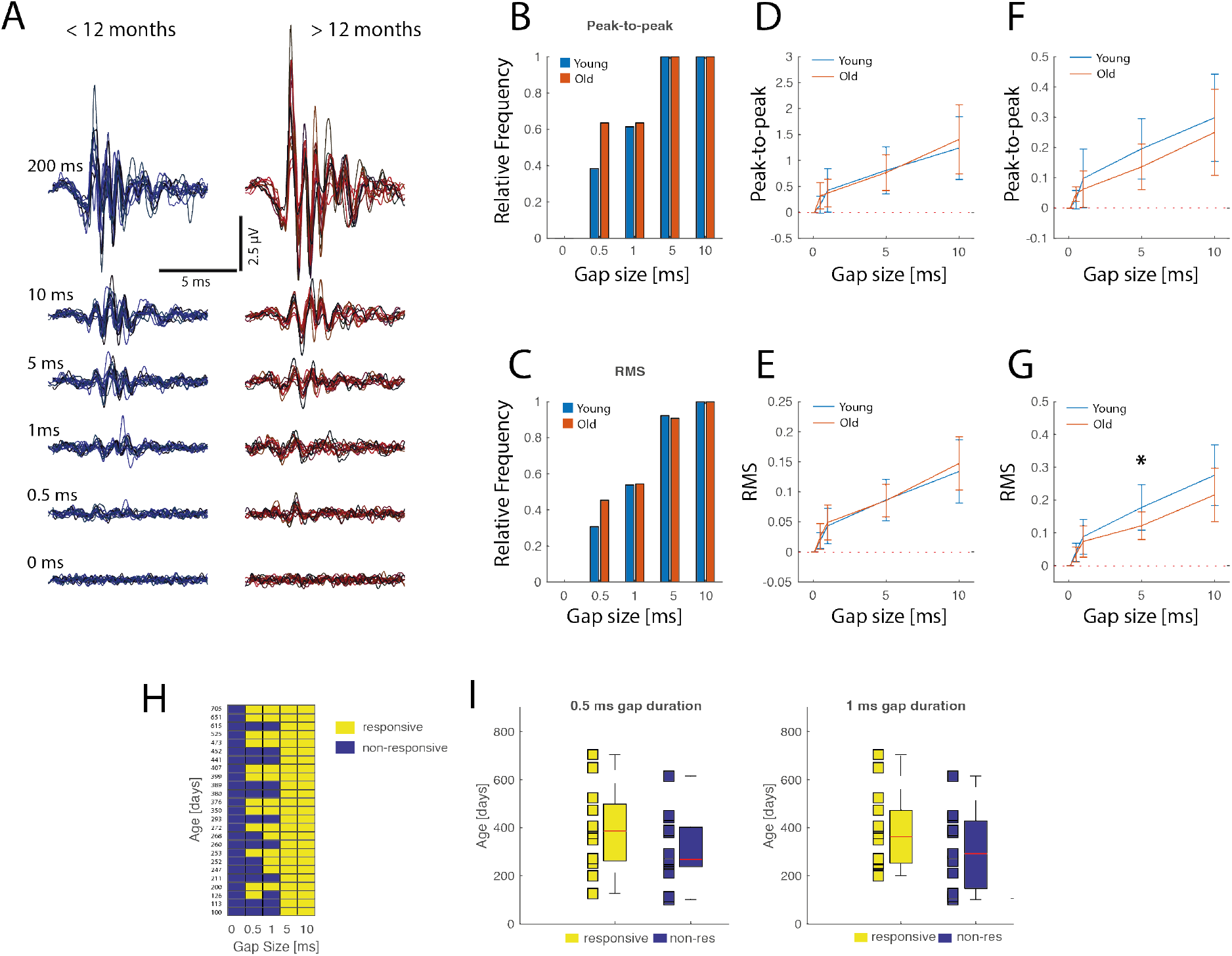
Auditory brainstem gap sensitivity remains robust over lifetime in normal hearing mice. A) Comparison of gap ABR between young (left, blue, n=13) and old (right, red, n=12) mice for onset and decreasing gap durations. B) Relative frequency of mice in both age groups showing a significant response (see methods) to gap durations measured for ABR peak-to-peak amplitude and C) RMS of the ABR. D) Change in peak-to-peak amplitude in dependence of gap duration for young and old mice corrected to baseline. E) RMS change in dependence of gap duration for young and old mice, corrected to baseline. F) Recovery ratio of Gap/Onset response amplitude change in dependence of gap duration for young and old mice, corrected to baseline. G) Recovery ratio of Gap/Onset response RMS change in dependence of gap duration for young and old mice, corrected to baseline. Error bars denote standard deviations. H) Overview of significant ABR responses (yellow) to different gap durations for all mice ordered by age (oldest top, youngest bottom). Significant responses are shown in yellow, and non-significant responses in blue. I) Comparison of ages for mice showing significant responses versus mice showing no responses for 0.5 ms gap duration (left). Comparison of ages for mice showing significant responses versus mice showing no responses for 1 ms gap duration (right).

The gap response recovery ratio reflects the relative change of the gap response in relation to the onset response of the beginning noise burst. Aging decreased the gap recovery ratio of the amplitude **(Figure 3F)** and RMS of the response **(Figure 3G)**. Thus, gap responses became comparatively smaller in relation to the onset response in old mice. For amplitude, two-way ANOVA did not show a significant interaction for the normalized peak-peak amplitude to onset response (F(4,115)=0.75 p=0.57). Simple main effects analysis confirmed that gap duration had a statistically significant effect on response amplitude (F(4,115)=42.18 p=0), and age showed a clear trend, however did not reach significance (F(1,115)=3.36 p=0.07). Two-way ANOVA showed also no interaction for normalized RMS to onset response F(4,115)=1.2 p=0.3. Simple main effects analysis confirmed that gap duration had a statistically significant effect on response amplitude (F(4,115)=82.31 p=0), and also age did have a statistically significant effect (F(1,115)=10.49 p=0.002). Post-hoc analysis revealed a significant difference at 5 ms (p=0.01).

Analysis of individual gap detection sensitivity over age confirmed significant responses to gap durations of 5 ms and 10 ms in all mice of our sample of CBA/J mice **(Figure 3H)**. More importantly, no age difference was present for a gap duration of 0.5 ms and 1 ms **(Figure 3I)**. We **c**ompared the age of mice exhibiting significant responses (responders) at 0.5 ms and 1 ms gaps to the age of mice who did not show responses at these gap durations (nonresponders). A two-sample t-test showed no significant difference in the age between significant responders (394 ± 173 days; mean/SD) and non-responders (309 ± 143 days; mean/SD); (t(23) = 1.34, p = .19) responses at 0.5 ms (**Figure 3I, left)**. Similarly, there was also no significant age difference between significant responders (384 ± 156 days; mean/SD) and non-responders for gap durations at 1 ms (307 ± 164 days; mean/SD); (t(23) = 1.19, p = .24) (**Figure 3I, right)**.

### 3.4 Multiunit gap sensitivity in the auditory midbrain is only minimally affected in aged mice with normal hearing

Next, we tested changes in IC multiunit activity of old mice with normal hearing. Multiunit gap responses in the IC of the youngest (<300 days, n= 10, 216±71 days; mean/std) and oldest (>=615 days, n=5, 657±42 days; mean/std) mice were comparable down to 1 ms gap duration **(Figure 4A)**, with slight differences in gap response strength at very short gaps **(Figure 4B)**. Responsiveness to gaps showed no clear pattern to reduced gap sensitivity at higher ages for multiunit population activity in the IC **(Figure 4C)**. Two-way ANOVA did not show a significant interaction for the *normalized peak response* to onset response (F(5,138)=0.08 p=0.99). Simple main effects analysis confirmed that gap duration had a statistically significant effect on the response amplitude (F(5,138)=14.24 p=0), but age did not have a statistically significant effect (F(1,138)=1.9 p=0.17). Further, analyzing the relative multiunit increase to baseline **(Figure 4D)**, two-way ANOVA did not show a significant interaction for the *relative amplitude increase to baseline* (F(5,138)=0.08 p=0.99). Simple main effects analysis confirmed that gap duration had a statistically significant effect on response amplitude (F(5,138)=15.42 p=0), but age did not have a statistically significant effect (F(1,138)=0.64 p=0.42). Analysis of individual gap detection sensitivity over age confirmed significant multiunit responses to gap durations of 5 ms and 10 ms in all mice of our sample of CBA/J mice **(Figure 4F)**, and most showed significant responses to 2 ms gaps. We **c**ompared the age of mice exhibiting significant multiunit responses (responders) at 1 ms gaps to the age of mice who did not show responses at these gap durations (non-responders). A two-sample t-test showed no significant difference in the age between significant responders (422 ± 187 days; mean/SD) and non-responders (355 ± 165 days; mean/SD); (t(23) = −0.94, p = .35) (**Figure 4F, right)**.

**Fig. 4.**
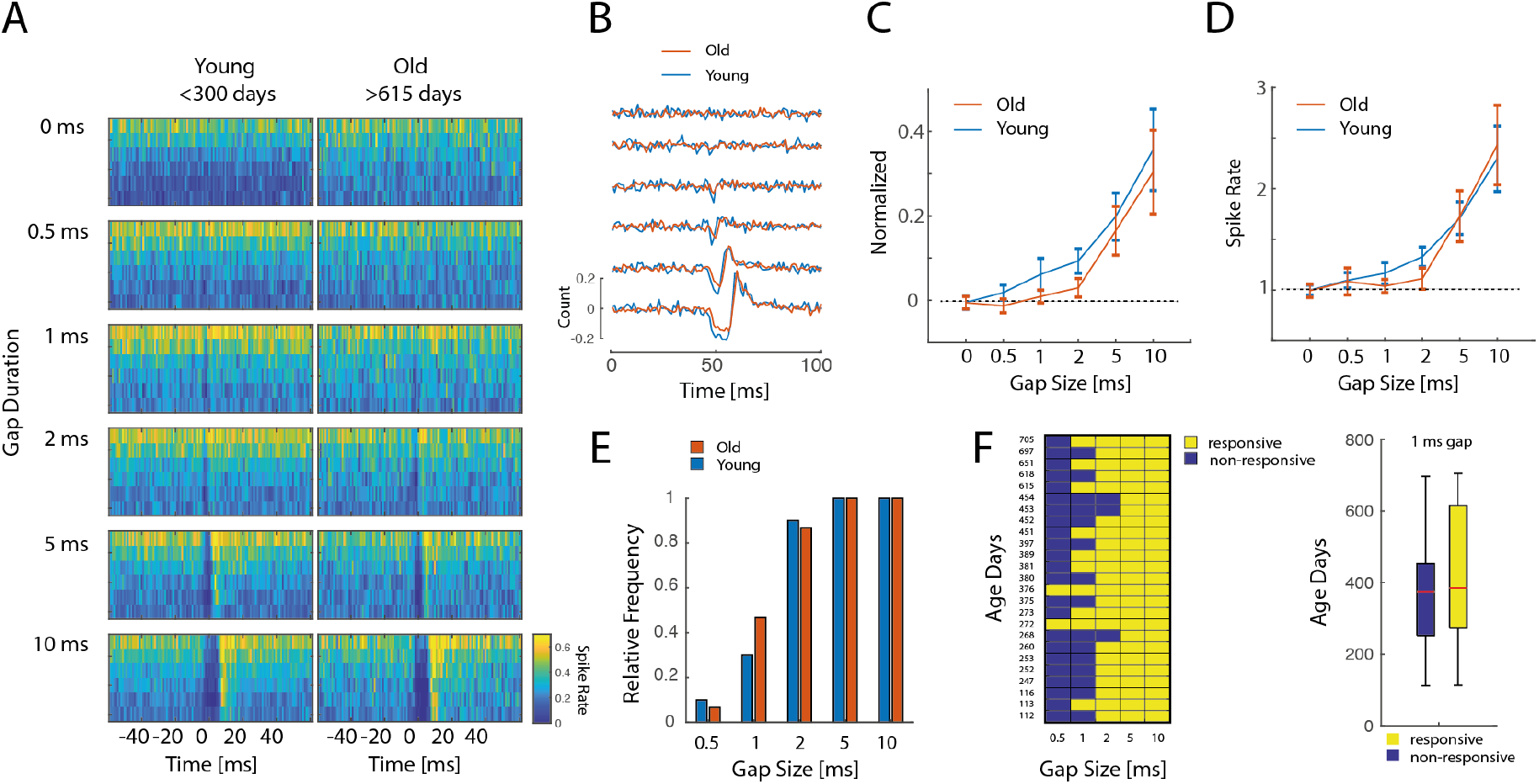
Multiunit gap sensitivity in the auditory midbrain is only minimally affected in aged mice with normal hearing. A) Comparison of multiunit gap responses in the IC between the youngest (<300 days, n= 10, 216±71 days; mean/std) and the oldest (>=615 days, n=5, 657±42 days; mean/std) mice. Color-coded rate plots show very similar activity for decreasing gap durations between young and old. Y-axis for each plot contains the 6 strongest electrode positions in IC. B) Grand average over mean-normalized responses over young mice (n= 10, 216±71 days; mean/std, blue) and old mice (n=15, 492±125 days; mean/std, red) and positions for decreasing gap durations. C) Change in normalized gap response strength for different gap duration corrected for young (n= 10, 216±71 days; mean/std) and old (n=15, 492±125 days; mean/std) mice. Error bars denote the standard error of the mean. D) Relative change in multiunit response to baseline in dependence of gap duration for young and old mice. Error bars denote the standard error of the mean. E) Relative frequencies of gap sensitivity in young and old mice. The bar plot shows that both groups are effectively similar. F) Gap sensitivity for each mouse for increasing age. No clear pattern related to age is visible. On the right boxplots, a comparison of responsive to non-responsive sites shows no differences in age.

### 3.5 Auditory midbrain activity of complex rhythmic sound is delayed (but robust) in aged mice with normal hearing

Next, we compared the reliability of multiunit representations of temporally complex rhythmic sounds between young and old mice **(Figure 5A**). The stimulus contained a complex rhythmic sound pattern (see methods) and multiunit activity in the IC was recorded during the signal presentation in free-field. The multiunit responses were robust in both young (n=10; age range: 112-273 days; 216±71 days; mean/std) and old mice (n=7; age range: 450-705 days; 612±85 days; mean/std) and closely represented the amplitude envelope of the signal **(Figure 5A, blue and red trace)**. Importantly, multiunit responses in old mice were delayed in comparison to responses in young mice, reflected by a shift in the crosscorrelation peak between mean multiunit responses of both groups **(Figure 5B)**, although the correlation was generally high. Pair-wise comparison between individual mice in the young and old group revealed a mean delay of the cross-correlogram peak of 1.2 ± 2.7 ms (mean/SD) in old mice (one-sample t-test, t(69)=3.67, p<0.001) **(Figure 5C)**. In addition, we compared the highest evoked peaks (n=52, threshold > 0.07 spikes/ms) between the age groups during sound presentation **(Figure 5D)**. These also showed a significant onset delay in aged animals, with additional differences in the later phase of the response (two-sample t-test with Bonferroni-Holm correction for multiple comparisons, p<0.05). Next, we calculated pair-wise cross-correlation of multiunit representations within young (n = 10; 45 pairs), within old (n = 7; 21 pairs), and between old and young mice (70 pairs) **(Figure 5E)**. Here we observed no differences in the mean correlation strength between multiunit representations of the groups (one-way ANOVA, F(2,133)=2.93 p=0.056), which supports the stability of IC multiunit responses across age.

**Fig. 5.**
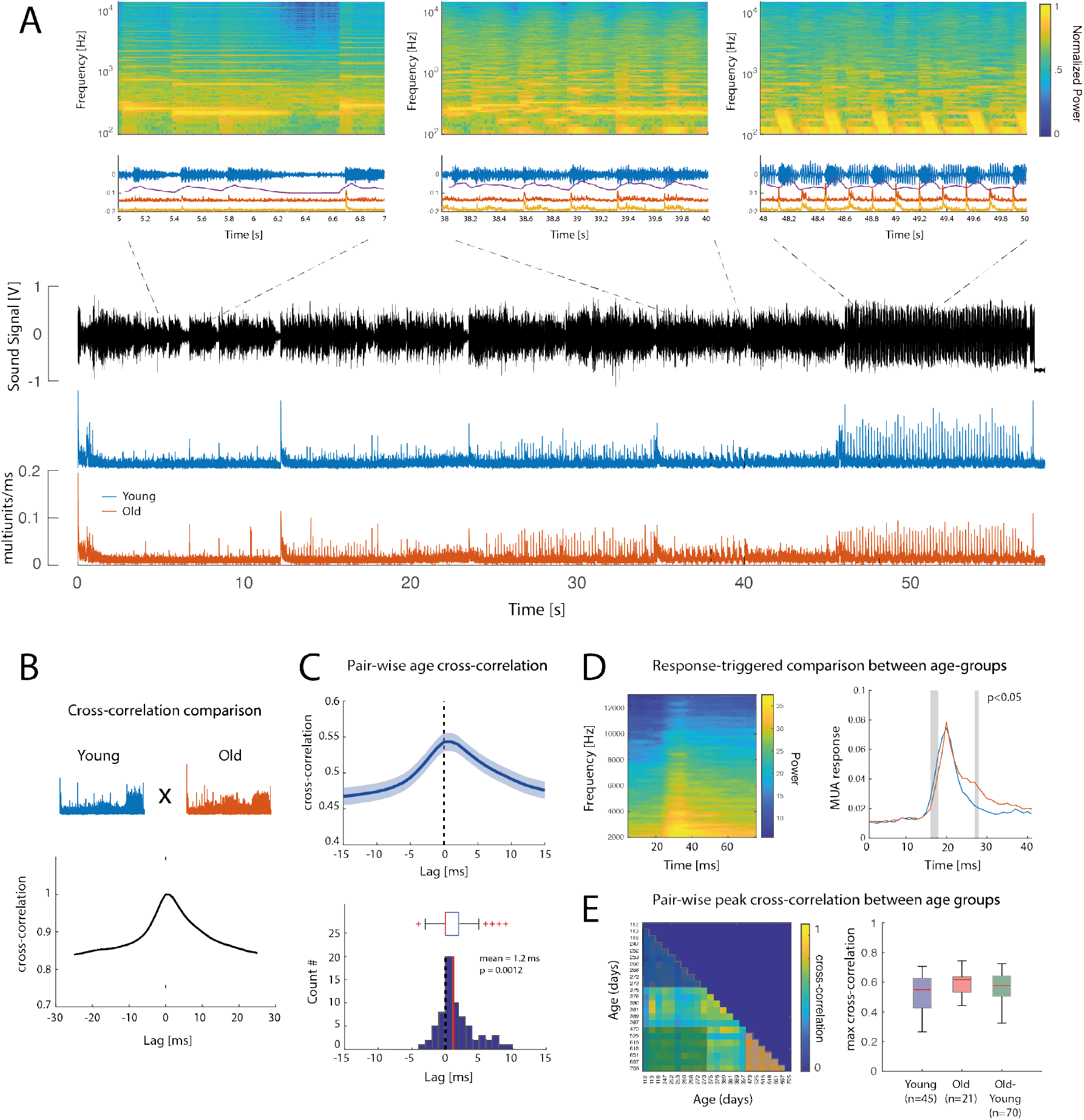
Auditory midbrain activity of complex rhythmic sound is delayed (but robust) in aged mice with normal hearing. A) Complex sound signal (black) with mean evoked multiunit activity in the auditory midbrain of young (blue) and old (red) mice over a duration of 55 s. Above enlarged 2 seconds intervals with parallel evoked multiunit activity and spectrogram. Note that spectrograms are slightly shifted in time (50 ms) in relation to the time signal, because of the moving-windowing. B) Cross-correlogram of sound-evoked multiunit activity of young (blue) and old (red) mice. Shown is the normalized cross-correlogram between the total averaged activity for each group (lower panel). C) Pair-wise correlation between the young and old age group (top). Histogram of lag distribution of the pair-wise correlation. D) Reverse-correlated spectrogram for the threshold-triggered multiunit activity for peaks above 0.07 spikes/ms (right). Comparison of response-triggered responses between young and old mice. Gray bars denote time intervals with significant differences between signals (p<0.05). E). Pair-wise cross-correlation matrix between all mice. Boxplots show within-groups (young, old) and cross-groups (young vs. old) comparisons.

### 3.6. Stability of auditory midbrain representations of complex sound is not reduced in old mice with normal hearing

Last, we tested the effects of aging on IC multiunit representation of 24 complex speech-like stimuli between young **(n=7; 260**±10 days; mean/SD) and old mice (n=6; **575**±127 days; mean/SD) **(Figure 6)**. These consisted of different short ‘words’, which contained no frequencies above 15 kHz to avoid age-related effects on mouse high-frequency hearing. This had the advantage to emphasize the envelope complexity instead of spectral properties and avoid the effects of high-frequency loss and peripheral effects. The ‘words’ produced distinct spatio-temporal representations in the IC **(Figure 6 A)**. The representations remained similar in very old mice in comparison to those in young mice **(Figure 6 A, B)**. To quantify this, we calculated pair-wise cross-correlations between individual mice of each age for each word **(Figure 6C)**. We then tested for differences in cross-correlation between young and old mice. There was no significant difference between young (n=7; 21 pairs), old (n=6; 15 pairs), and young-old (n=7×6; 42 pairs) cross-correlation respectively (one-way ANOVA, F(2,75)=0.17 p=0.84) **(Figure 6D, E)**. Last, we tested direct cross-correlation between the sound envelope and the population multiunit response for each word **(Figure 6F)**. Multiunit correlations were in general similar between young and old mice. Central aging is assumed to degrade such temporal representations but contrary to this expectation this effect was not found. Rather a trend towards better correlation in aged mice between multiunit response and ‘words’ was observed. When arranged in increasing order for the cross-correlation values, only four words showed non-overlapping bootstrapped confidence intervals and significant differences between age groups (subsequent pair-wise t-test, p<0.05) **(Figure 6F)**. Importantly, pooled comparison for all words for the two groups did not reveal a decline in general sound representation in the old group (two-sample t-test, t(11)=-2.07, p=0.062), however showed a trend towards an increase in representation precision in old mice **(Figure 6G)**.

**Fig. 6.**
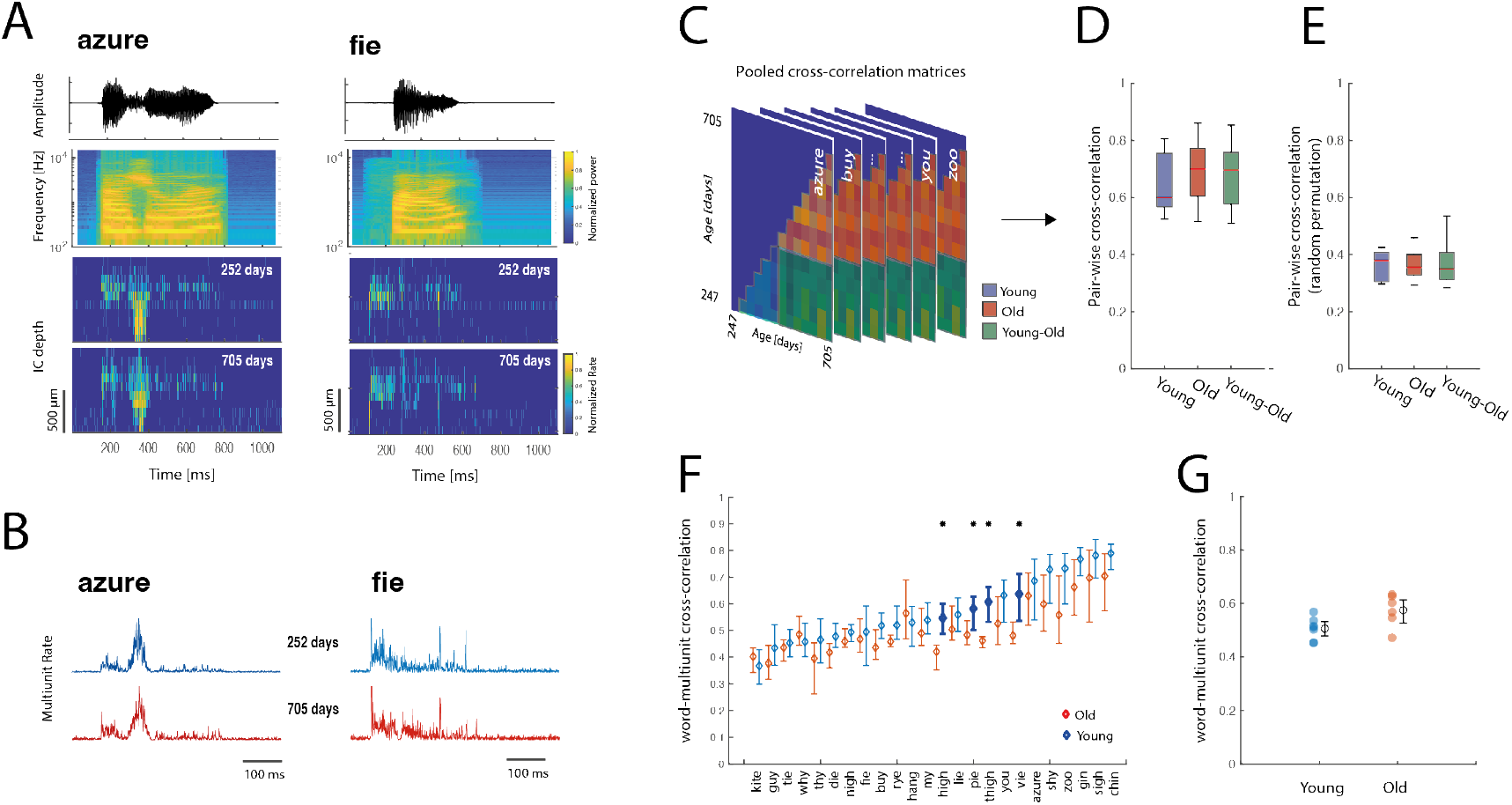
Stability of auditory midbrain representations of complex sound is not reduced in old mice with normal hearing. A) Top row: Soundwave of a female voice speaking the words ‘azure’ and ‘fie”. Below spectrograms are shown of the soundwaves above. Two bottom rows: Depth profile of multiunit responses in the IC to the sounds above in a young (256 days) and old mouse (705 days). B) Examples of depth-averaged multiunit responses in A) for the two words. C) Pairwise cross-correlation matrices between the neural activity of all individual mice for each word. D) Boxplots show pooled correlation within age groups (young, old) and across age groups (young vs. old) comparisons for all words (n=23). E) Control correlations for same analysis as in D) with randomly permutated neural activity. F) Mean word-multiunit response correlations in young (n=7; range 247-273 days) and old mice (n=6; range 400-705 days) for each word. Mean correlations are sorted in ascending order with bootstrapped confidence intervals for each word. G) Pooled correlations across words for each age group (p > 0.05).

## 4. Discussion

We demonstrated that age-related slowing of auditory brainstem conduction time – in the sub-millisecond range-did not have a notable effect on temporal resolution and complex sound representations in the auditory midbrain of old mice when peripheral hearing remained normal in aging.

### 4.1 Limited effect of age-related delays on temporal resolution and complex sound representations in the auditory midbrain

Age-related slowing of central conduction time in two-year-old normal hearing CBA/J mice summed up to around 700 μs in the ABR I-V interval. Similar sub-millisecond age-related shifts in ABR interpeak intervals have been described in aging mice, rats, and guinea pigs (Cooper et al., 1990; Backoff and Caspary, 1994; Ingham et al., 1998; Hong et al., 2007; Chen et al., 2010; Mock et al., 2011; Beltrame et al., 2021) and humans (Rowe, 1978; Kjaer, 1980; Jerger and Hall, 1980; Maurizi et al., 1982; Harrison and Buchwald, 1982; Allison et al., 1983, 1984; Kelly-Ballweber and Dobie, 1984; Chu, 1985; Rosenhall et al., 1986; Stürzebecher and Werbs, 1987; Elberling and Parbo, 1987; Jerger and Johnson, 1988; Lenzi et al., 1989; Mitchell et al., 1989; Walton and Burkard, 2001; Burkard and Sims, 2002; Anderson and Karawani, 2020). In addition, we observed an age-related delay of IC multiunit onset latency at the level of the auditory midbrain around 1 ms and in the ABR I-P_0_ interval – with P_0_ reflecting an IC component in the mouse (Land et al., 2016; Duque et al., 2018). Latency changes in aging has also been described in the IC of guinea pigs (Dum, 1983) and mice (Simon et al., 2004). However, not all studies consistently find significant age-related slowing in the brainstem (Beagley and Sheldrake, 1978; McClelland and McCrea, 1979; Otto and McCandless, 1982; Jerger and Johnson, 1988), age-related central slowing in the midbrain appears to remain largely limited to the sub-millisecond, near millisecond range, in cases when it occurs.

The small shift of age-related slowing of conduction time did not lead to significant changes in the brainstem and auditory midbrain gap sensitivity in normal hearing aging mice. When controlled for peripheral hearing loss, gap detection thresholds in aging humans also remain at maximum limited to millisecond changes, with old subjects often exhibiting very good gap detection thresholds, challenging the idea of an automatic uniform decline during aging (Snell, 1997; Strouse et al., 1998; Walton et al., 1998; Harris et al., 2012; Palmer and Musiek, 2014; Hoover et al., 2015; Ozmeral et al., 2016; Sanju et al., 2017). In humans, gap thresholds shifts are limited to the millisecond range (Palmer and Musiek, 2014), with surprisingly precise ABR gap sensitivity in aged subjects (Poth et al., 2001). In mice, ABR gap detection thresholds remained as low as 2 ms in young and middle aged CBA/J mice (Williamson et al., 2015), with minor changes in gap recovery functions, similar to studies in aged gerbils (Boettcher et al., 1996; Hamann et al., 2004).

Such stability of temporal resolution in aging has been described in the IC and cortex of normal hearing mice (Osterhagen and Hildebrandt, 2018; Land and Kral, 2022), and age-related changes in central temporal acuity in the midbrain can often be explained by inherited peripheral effects (Land and Kral, 2022). Thus, the issue of confounding peripheral effects remains central, as existing studies on gap detection (Barsz et al., 2002; Walton et al., 2008; Ross et al., 2010; Shen, 2014) indicate an upper limit of pure age-related changes in the auditory midbrain if no other pathological states are present.

Regarding a role in perceptual deficits, the question is, whether small brainstem changes effectively deteriorate envelope representation of complex sound patterns occurring at slower time scales. For ‘pure’ central presbycusis independent of cochlear damage, small changes in conduction time and temporal resolution – when present – need to substantially affect ‘speech’ complex sound representations (Gates, 2012; Humes et al., 2012). On a neuronal level, we did not observe gross changes between young and old mice in midbrain neural sound envelope representations, besides a small temporal delay, reflecting ‘remarkable normalcy in aging IC activity in old CBA/J mice’ (Willott et al., 1988). Changes in simple measures such as temporal gap detection, and envelope following responses have been shown to be only a weak predictor of speech perception in aging (DeMetropolis et al., 2015; Hoover et al., 2015; Schoof and Rosen, 2016), and speech perception deficits can often more simply explained by spectral decline present in ‘clinically normal’ hearing elderly (Gelfand et al., 1985, 1986).

The observation that most elderly have in principle no problems in speech perception in quiet when hearing is normal (Schoof and Rosen, 2014) supports general stability of central processing in aging. Also, automatic central pathway deterioration in aging per se would limit the effectiveness of hearing aids. Thus, the general stability of auditory perception during the lifespan of individuals with normal peripheral hearing questions the notion of automatic age-related degradation of central auditory transmission.

### 4.2 Relation to other age-related factors influencing hearing problems in aging

Our results support the differentiation of auditory pathway stages with respect to central presbycusis. Although brainstem, auditory midbrain, and thalamus population processing often is stable in old animals with normal hearing (Lee et al., 2002; Mendelson and Lui, 2004; Osterhagen and Hildebrandt, 2018; Land and Kral, 2022), small effects may cumulate towards higher levels of the auditory system (Mendelson and Lui, 2004; Occelli et al., 2019; Quraishe et al., 2020), affecting the precision of cortical processing, which is critical for temporal and complex sound and processing (Nourski and Brugge, 2011; Nourski et al., 2021a, 2021b). Elongation of cortical potentials is present in aged mice (Mei et al., 2021), and middle latency and cortical evoked potentials in aged humans (Goodin et al., 1978; Dum, 1983; Allison et al., 1984; Martini et al., 1991; Gourévitch and Edeline, 2011; Price et al., 2017; Occelli et al., 2019). This could explain the central aging effects of processing (besides simple latency) on the cortical level (Weible et al., 2014; Occelli et al., 2019), and higher processing stages (Juarez-Salinas et al., 2010). This might be related to general age-related slowing, affecting perceptual processing and other sensory systems (Allison et al., 1984; Gilmore, 1995; Eckert, 2011). Cumulative slowing towards higher stages may also smear the transition towards higher ‘auditory-cognitive’ processing.

Regarding the relative contribution of central auditory pathway changes on perception in comparison to peripheral cochlear and cognitive aging (Lentz et al., 2022), it is undisputed that peripheral deficits play a pivotal indirect role (Ison et al., 2010). Perceptual resolution of human temporal acuity is established in the cochlea and maintained at progressively higher levels of the hearing pathway (Bidelman and Bhagat, 2017). It is a recognized problem that peripheral confounding factors in studies on age-related central decline (‘pure’ central effects), especially in psychophysical studies contribute to conflicting results of central aging (Humes et al., 2012). Aging effects on gap detection can be confounded by presentation level and undetected peripheral hearing loss, explaining why gaps at lower levels are affected in aging whereas higher levels are not (He et al., 1999; Allen et al., 2003). Further, non-auditory age-related cognitive factors affect gap sensitivity and central measures (Schneider and Hamstra, 1999; Harris et al., 2010; Ross et al., 2010), or are improved with musical training (Zendel and Alain, 2012). As cognitive decline often is a good predictor for age-related perception deficits (Füllgrabe et al., 2015; Lentz et al., 2022), the stability of IC representations speaks for only a small contribution in aging in relation to peripheral and cognitive decline.

In terms of effect size and functional relevance of age-related central changes on auditory perception, the existing evidence on midbrain gap detection or brainstem conduction speed indicates that the effects are at best small and/or highly variable. Studies may be often underpowered, which explains variability in findings, and increase the likelihood of false positives (Button et al., 2013), overemphasizing the impact of aging. Also, multiple factors and isolated small central effects still may add up from multiple causes (Götz et al., 2021). However, the impact of small neural changes on self-reported perception deficits in normal hearing elderly – independent of peripheral damage or cognitive factors - remains to be further explored.

## 5. Conclusion

Speech-like sound representations in the auditory midbrain remained stable despite age-related brainstem slowing in old normal hearing mice. This finding suggests the general stability of neural midbrain processing in aging mice. It points out the importance to assess the impact of small neuronal effects on perception, their possible cumulative effects on higher auditory processing centers, and understanding the relation of peripheral, cognitive, and central factors in auditory perception deficits in the aging brain. Most importantly it emphasizes the ultimate importance of peripheral input preservation in aging.

## Acknowledgments

We are grateful for technical and logistical lab support by Karl-Jürgen ‘Eddy’ Kühne and Daniela Kühne. This work was supported by grants from the DFG Priority Program: Ultrafast and Temporally Precise Information Processing: Normal and Dysfunctional Hearing (SPP 1608; 198624999), Cluster of Excellence (Exc 2177), and MEDEL (Innsbruck, Austria).

